# Diverging trends and drivers of Arctic flower production over space and time

**DOI:** 10.1101/2022.11.15.516565

**Authors:** Antoine Becker-Scarpitta, Laura H. Antão, Niels Martin Schmidt, F. Guillaume Blanchet, Elina Kaarlejärvi, Katrine Raundrup, Tomas Roslin

## Abstract

The Arctic is warming at an alarming rate. While changes in plant community composition and phenology have been extensively reported, the effects of climate change on reproduction remain poorly understood. We quantified multidecadal changes in flower density for nine tundra plant species at a low- and a high-arctic site in Greenland. We found substantial changes in flower density over time, but the temporal trends and drivers of flower density differed both between species and sites. Total flower density increased over time at the low-arctic site, whereas the high-arctic site showed no directional change. Within and between sites, the direction and rate of change differed among species, with varying effects of summer temperature, the temperature of the previous autumn and the timing of snowmelt. Finally, all species showed a strong trade-off in flower densities between successive years, suggesting an effective cost of reproduction. Overall, our results reveal region-and taxon-specific variation in the sensitivity and responses of co-occurring species to shared climatic drivers, and a clear cost of reproductive investment among arctic plants. The ultimate effects of further changes in climate may thus be decoupled between species and across space, with critical knock-on effects on plant species dynamics, food web structure and overall ecosystem functioning.

## Introduction

The Arctic is warming faster than the global average (IPCC 2021; Voosen 2021), with direct effects on permafrost, sea-and land-ice surface dynamics, on patterns in snow depth and snowmelt, and on ecosystem functioning (Box et al. 2019; Post et al. 2019). In this context, the arctic tundra biome offers an exceptional “natural laboratory” to test hypotheses related to the effects of climate change on ecological communities. Although extensive research has been conducted on shifts in the phenology of arctic plants and their traits in response to climate change (Panchen and Gorelick 2017; Prevéy et al. 2019, 2021; Bjorkman et al. 2018, 2020; Collins et al. 2021), the temporal dynamics of flower production (i.e., the reproductive output) remain less explored. For plants, sexual reproduction with flowers and seeds is a key strategy to maintain local genetic diversity and to disperse in space and time (Körner 2003). Nonetheless, despite their potential importance, changes in flower production in arctic plants have received only limited attention at the species level (Høye et al. 2007a; Semenchuk et al. 2013; Kelsey et al. 2021) and even less at the community-or site level. Much of this knowledge deficit may be due to a lack of appropriate long-term data. As a result, insights into how the long-term effects of climatic conditions and population dynamics may modulate flower production from year to year are largely lacking.

From the perspective of plant-associated taxa, flowers are an essential resource. For example, many arctic insects depend on flowers for food, providing pollination services for plants in return (Kevan 1972). Thus, changes in the dynamics of flower production or in the composition of plant communities can profoundly affect local ecological networks and ecosystem functioning (Schmidt et al. 2016; Tiusanen et al. 2020). Given the recent and rapid environmental changes in the Arctic (Robinson 2022), a better understanding of the mechanisms determining flower production is fundamental for quantifying and predicting population, community, and ecosystem dynamics of both plants and their associated taxa.

Predicting how individual plant species and communities respond to climatic changes is a key challenge. For instance, there is high variation in the rate and direction of phenological change responses across both species and regions (Thackeray et al. 2016; Delgado et al. 2020; Roslin et al. 2021), including in the tundra biome (Prevéy et al. 2019; Bjorkman et al. 2020; Kelsey et al. 2021). In addition, species respond to multiple environmental drivers. Snow conditions have been identified as a key driver of Arctic vegetation dynamics (Callaghan et al. 2011; Niittynen and Luoto 2018; Niittynen et al. 2018; Happonen et al. 2019), impacting both microclimatic conditions (e.g., protecting plants from dry and cold air in winter and early spring) and the start of the growing season (Inouye 2008; Bokhorst et al. 2012, 2016; Cooper 2014; Post et al. 2019). Thus, the timing of arctic plant flowering seems strongly associated with snowmelt dates (Semenchuk et al. 2013). In addition, higher temperatures and longer growing seasons have been found to positively affect flower abundance (Semenchuk et al. 2013; Prevéy et al. 2021), while low temperatures early in the season might cause flower abortion, thus decreasing reproductive success (Körner 2003; Høye et al. 2007a; Inouye 2008).

Importantly, the phenology of early-flowering species seems to be mainly determined by photoperiod and snowmelt dates, while late-flowering species are mostly driven by heat accumulation over the spring and summer (Molau 1993; Wipf 2010). From these patterns, direct parallels can be drawn to flower production, since phenology, growth and flower production have been found to respond to the same environmental variables (Krab et al. 2018; Kelsey et al. 2021). However, it remains unclear how different climatic variables impact flower production.

In addition to abiotic impacts, long-term flower production can be affected by strategies for interannual resource allocation by the plant itself (Obeso 2002; Høye et al. 2007a, b). In the Arctic, most perennial species accumulate resources over the growing season to produce flower buds in late summer or early autumn. This “bud pre-formation strategy” amounts to investing resources for the following flowering season. Early-flowering species generally benefit from well-differentiated floral buds initiated in the previous year, whereas late-flowering species require bud maturation or even full bud formation during the growing season (Körner 2003). The production of flowers in a given year may thus be affected by climatic conditions of the current year, as well as by climate and resource allocation for reproduction in the previous year. Such temporal dependence between years - i.e., the trade-off between flowering in a given year *vs* the next - may offset resources for flowering in the current year (Obeso 2002; Høye et al. 2007a; Semenchuk et al. 2013). While warming appears to result in a contraction of the community-level flowering season in tundra ecosystems (Prevéy et al. 2019), the same warming increases the length of the growing season, which may in turn increase biomass accumulation and thus the resources available for flower production (Chapin and Shaver 1996; Lund et al. 2010; Lyngstad et al. 2017). Given these trade-offs and dependencies, it remains unclear how warming affects species-specific investment in reproductive structures, and whether this is potentially reflected in asymmetric responses among co-occurring plant species.

Compounding spatiotemporal patterns in plant flowering is the fact that both current and projected climatic conditions vary substantially between different regions of the Arctic (Abermann et al. 2017; Prevéy et al. 2017; Nabe-Nielsen et al. 2017; Bhatt et al. 2021). As different aspects of the environment may becoming limiting under different climatic regimes, it is thus crucial to compare the impact of individual drivers on flower production across multiple sites and under different conditions.

Here, we assessed temporal changes and underlying drivers of flower production over two decades at two sites located ca 1000 km apart, on different sides of Greenland. For this purpose, we use time series of flower density for nine typical arctic tundra species. Given the spatial heterogeneity of climate change and vegetation composition, we tested the following hypotheses:

- **(H1) climatic trends over the last few decades have differed between sites**, showing different rates of change between sites for different climate variables.
- **(H2) total annual flower production increased over time**, with a stronger increase at sites experiencing faster warming.
- **(H3) plant species showed different temporal trends in flower production, underpinned by different climatic variables**. Assuming that patterns in flower production mimic those reported for phenology, we expect some species to take advantage of late snowmelt dates and warm previous year autumns, while others would benefit from early snowmelt and long and warm growing seasons.
- **(H4) flower production in a given year showed a trade-off against flower production in the next.** Given limited resources, we expect that higher investment in reproduction in one year will translate into lower investment in the following year.

## Material and Methods

Flower abundance and climatic data were extracted from *The Greenland Ecosystem Monitoring Database* (GEM, https://data.g-e-m.dk, extracted in 2022).

### Study sites

To compare trends and drivers of flower production between regions characterised by different climatic conditions and trends, we focused on two regions in Greenland for which we had access to decade-long flower abundance data: the Kobbefjord Research Station, located in the low Arctic (Southwest Greenland, 64° 08’ N, 51° 23’ W), and the Zackenberg Research Station located in the high Arctic (Northeast Greenland, 74° 29’ N, 21° 34’ W, Fig 1).

**Figure 1:**
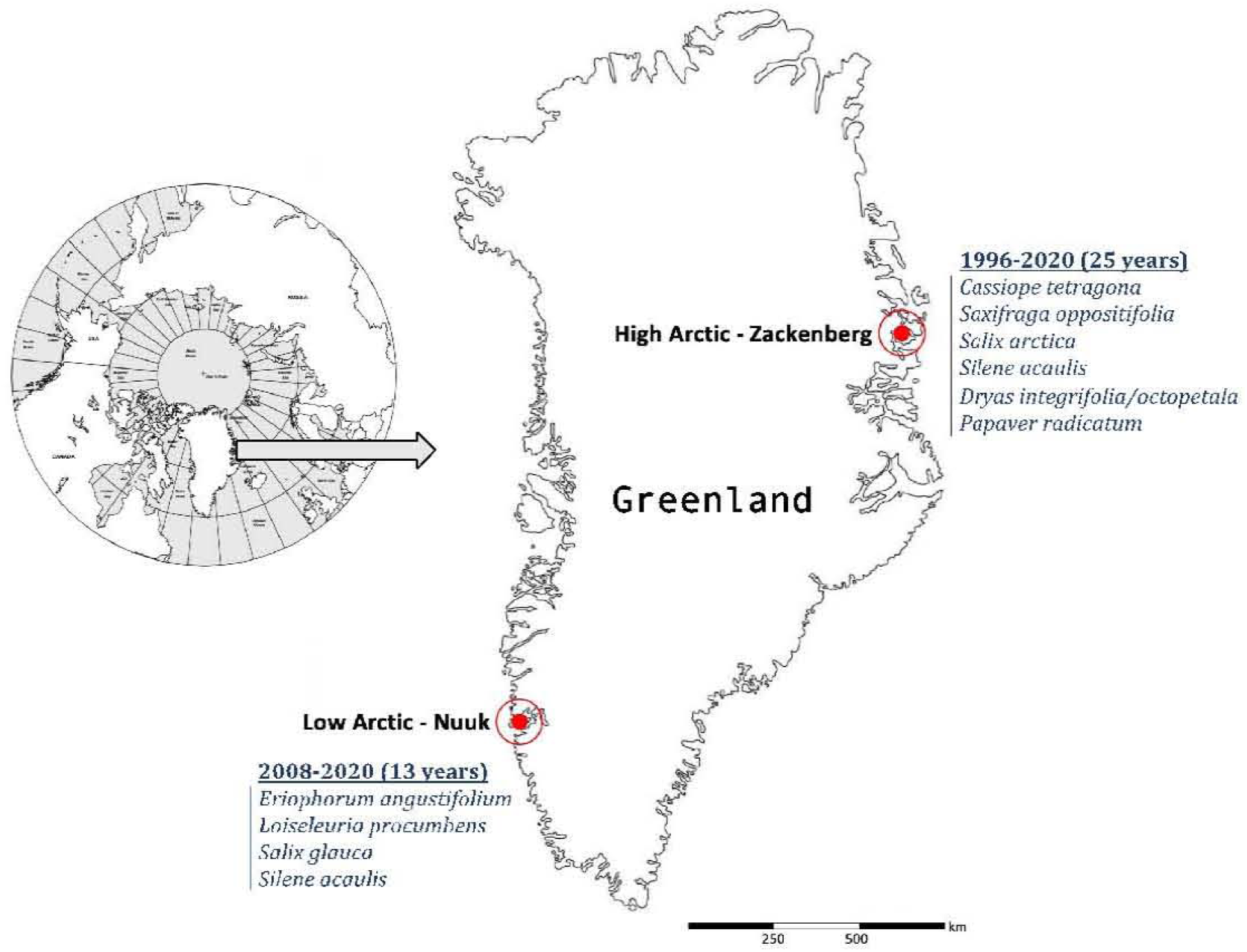
Map of the study sites, length of the local time series and list of species surveyed locally for flower abundances.

At the low-arctic site, for the 2007-2020 period, the mean annual temperature was - 0.06°C. July was the warmest month with an average temperature of 10.6 °C (ranging from −0.9 to 22.3°C), while the coldest was February with an average of −8.6°C (ranging from −30.2 to 9.2°C). The region is characterized by spatially discontinuous permafrost, and annual mean precipitation (rain and snow combined) of ~838 mm. Terrestrial ecosystems in this area are diverse and include vegetation types typical of the low-arctic region: fen, copse, grassland and dwarf shrub heath dominated by *Betula nana* L., *Vaccinium uliginosum* L., *Empetrum sp., Salix sp.* and *Cassiope tetragona* (L.) D. Don (CAVM Team 2003; Walker et al. 2005). At the high-arctic site, for the same period, the mean annual temperature was −8.6 °C. July was the warmest month with an average temperature of 6.8 °C (ranging from −1.9 to 19.6°C), while the coldest is March with an average of −20.9°C (ranging from −38.4 to 2°C). In this region, permafrost is spatially continuous and total annual precipitation amounts to ca 260 mm. As in the low-arctic site, various tundra vegetation types can be found, from barren ground to fen and heaths dominated by dwarf-shrub species such as *Cassiope tetragona* (L.) D. Don and *Salix arctica* Pall.

### Flower abundance data

The data consist of annual plot-level time series, where the total number of flowers was counted at the peak of the flowering season for nine plant species (Appendix S1). At the start of the monitoring, species-specific plots were established to cover the full range of species’ ecological niches (details of the sampling design are given in Appendix S2). In the high Arctic, six plant species were then surveyed over 25 years (1996-2020), while in the low Arctic, four species were surveyed over 13 years (2008-2020). All the species included in the monitoring program are perennial: *Saxifraga oppositifolia* L.; *Loiseleuria procumbens* (L.) Desvaux; *Salix arctica* L.; *Salix glauca* L.; *Eriophorum angustifolium* Honck.; *Papaver radicatum* Rottb.; *Cassiope tetragona* (L.) D. Don; *Silene acaulis* (L.) Jacq; and the hybrid *Dryas integrifolia/octopetala* (for further details on species, see Appendix S3).

Each plot was divided into four identical sections: A, B, C, and D, and the subsections were visited once a year, on the same day to quantify the total number of flowering structures. Subplots were pooled at the plot level since the count of some subplots might be very low. The exact date of the survey varied between species, plots, and years depending on climatic conditions. Reproductive structures counted include *flowers, flower buds* (not sexed for *Salix),* and *senescent flowers* (Schmidt, Hansen, et al. 2019a, Raundrup, Olsen, et al. 2020). For *Salix* species, we counted catkins, and the total flower number was recorded separately for males and females in each plot. Since *buds* are unsexed flowers, we split them into “male” or “female” based on the average long-term observed sex ratio of the *Salix* flowers for each plot independently.

To control for differences in plot area (ranging from 1 to 300m^2^), we calculated flower density per plot for each species by counting the number of flowers recorded in a plot within a given year and dividing it by plot area. For the site-level annual flowering density analysis (see SiteFD below), we summed the annual flower abundance of all species and divided them by the sum of all individual plot areas in each site. For the species-level flowering density analysis (see SpeciesFD below), the plot-level flower densities were log_e_+1 transformed.

### Environmental drivers

We extracted local climatic data for each of the sites. Specifically, we used air temperature (°C) and accumulated precipitation (mm) as recorded at the site level, thus resulting in a single value per year for all plots in each site. Both variables were recorded on average every 30 min at Nuuk, and every 60 min at Zackenberg. As such, we averaged all readings by day and by month for each year. We calculated the average seasonal temperature and precipitation for summer (June-August) and autumn (September-November), as well as annual mean temperature and precipitation values, to estimate long-term changes in each site. The period covers the years 2007-2020 for the low-arctic site and the years 1995-2020 for the high-arctic site.

For assessing snowmelt dates, we extracted the yearly measures recorded at the plot level, which are based on the visual estimate of snow cover made in the field 1-3 times every week for each plot section. We used linear interpolation to predict the estimated day of the year when a plot (i.e. all sections combined) reached 50% snow cover. Note that for some cases, the day of the year of 50% snow cover could not be estimated, as due to late arrival by the monitoring team to the field sites.

### Statistical analysis

Our analyses focused on resolving four imprints of environmental change: (i) the temporal trends in climatic conditions at each site, ii) the temporal trend in total annual flower density at the site level, (iii) the temporal trends in annual flower density at the species level, and (iv) the effects of the previous year flower density, as well as temperature and snowmelt date on flower density inter-annual dynamics.

First, to test for differences in recent climate change between the high-and low-arctic sites, we used linear models to quantify temporal trends in annual and seasonal mean temperature and precipitation, as well as snowmelt dates.

Second, to quantify the direction and magnitude of the temporal changes in the site-level total flower density (SiteFD, eq. 1), we modelled the flower densities as a function of year (Y, continuous variable), fitting a separate linear model for each site.

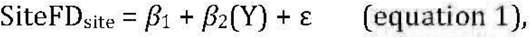

where *β_1-2_* are estimated parameters, and ε defines the model’s residuals (assumed to be independent and normally distributed). Following H1, we expect a significant increase in flower density over time, i.e., *β*_2_ >0.

Third, to quantify the direction and magnitude of the temporal change in species-level annual flower density (SpeciesFD, eq. 2), we modelled the log_e_+1 transformed flower densities as a function of the two-way interaction between species (S; categorical variable) and year (Y, continuous variable). We fitted a separate linear mixed model for each site.

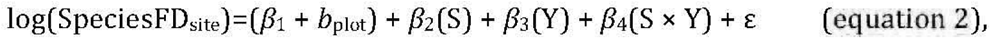

where *β_1-4_* are estimated parameters, *b*_plot_ is a random intercept capturing the effect of plot, and ε is the model’s residuals (assumed to be independent and normally distributed). Note that since S is a categorical variable with as many levels as there are species in each site, *β_2_* and *β_4_* are actually composed of multiple estimated parameters.

Following H2, we expect *β_4_* to be significant for each species, i.e., species to exhibit significantly different temporal trends.

Fourth, to disentangle the drivers of year-to-year variation in species’ annual flower densities (SpeciesFD, eq. 3), we modelled the interaction between species (S) and environmental drivers, i.e., average summer temperature of the current year (ST), snowmelt day of the year (SM), and previous year autumn temperature (PAT) for each site separately. We further accounted for temporal dependence (TD) in flower densities by including the previous year’s flower density (also loge+1 transformed). The different terms in eq. 3 aim at capturing the multiple climatic effects we aimed to test, specifically: ST was used to capture the temperature effect during the growing season; SM was used to capture the effect of the start of the growing season; PAT was used to capture the effect of resource accumulation at the end of the previous growing season, and TD was used to capture the effect of reproduction cost from the previous year. Since flower densities were modelled on a log-log scale, equal flower densities in consecutive years is conveyed by a regression coefficient of 1, whereas a coefficient <1 implies that high densities in a year translate into lower densities the next year. Thus, we tested TD for significant deviations from 1 (following Forchhammer et al. 2008). Technically, this was carried out by adding an offset of 1 to TD across species in the model. In this respect, we estimated 1-*β*_4_ and not *β*_4_ in the model. Thus, estimating *β*_4_>1 indicates more flowers are produced at time *t* than at time *t*-1; *β*_4_=1 indicates the same number of flowers are produced from year to year, and *β*_4_<1 means fewer flowers are produced at time *t* compared to time *t*-1. In other words, in the context of our study, we interpreted *β*_4_ as representing different types of temporal dependence (i.e., over-or under-compensation). To allow comparison between species and sites, summer temperature

(ST), snowmelt day (SM), and the previous autumn temperature (PAT) were all standardized to a mean of 0 and a standard deviation of 1.

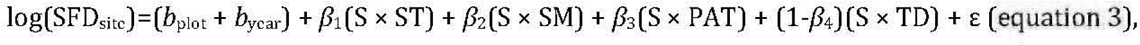

In equation 3, as for equation 2, *β_1-4_* are vectors of estimated parameters associated with each species, and ε defines the model’s residuals (assumed to be independent and normally distributed). In addition, *b*_plot_ and *b*_year_ are random intercepts accounting for the crossed effect of plot within years - i.e., they estimate variation resulting from the spatial and temporal structure of the experimental design. It is important to note that contrary to the other parameters because the parameters in *β*_4_ were tested against 1 (and not 0), the p-values associated with *β*_4_ needed to be calculated according to this particularity of the model and was estimated with the function *pt* within the “stats” package.

All statistical analyses were conducted in R (R Core Team, 2022). Linear mixed effect models were fitted with the function *lmer* within the ‘lme4’ package (v.1.1-14, Bates et al. 2015).

## Results

### Climatic change

In the low-arctic site, there was no statistically detectable trend in the average annual, summer and autumn temperatures over the 12 years of study (Appendix S4, Fig. S4a & S4b). However, we detected a decrease in the average annual precipitation (Appendix S4, Fig. S4a). By contrast, in the high-arctic site, annual, summer and autumn temperatures and precipitation increased over the last 25 years (Appendix S4, Fig. S4a & S4b). The snowmelt day of the year did not change in the low Arctic, while in the high

Arctic, snow tended to melt earlier in all sampling plots (with statistically supported trends in *Cassiope, Dryas, Papaver* and *Silene* plots (Appendix S4, Fig. S4c).

### Total flower production

The average site-level annual flower density at the high Arctic is much lower compared to the low-arctic site (Fig. 2). We detected an increase in the site-level annual flower density at the low-arctic site (β_2_=1.82 flowers/m^2^/year, CI[0.35-3.28], p=0.02, R^2^adj=0.35), while no change was observed for the high-arctic site (β_2_=−0.11, CI[−0.27-0.05], p=0.17, Fig. 2).

**Figure 2:**
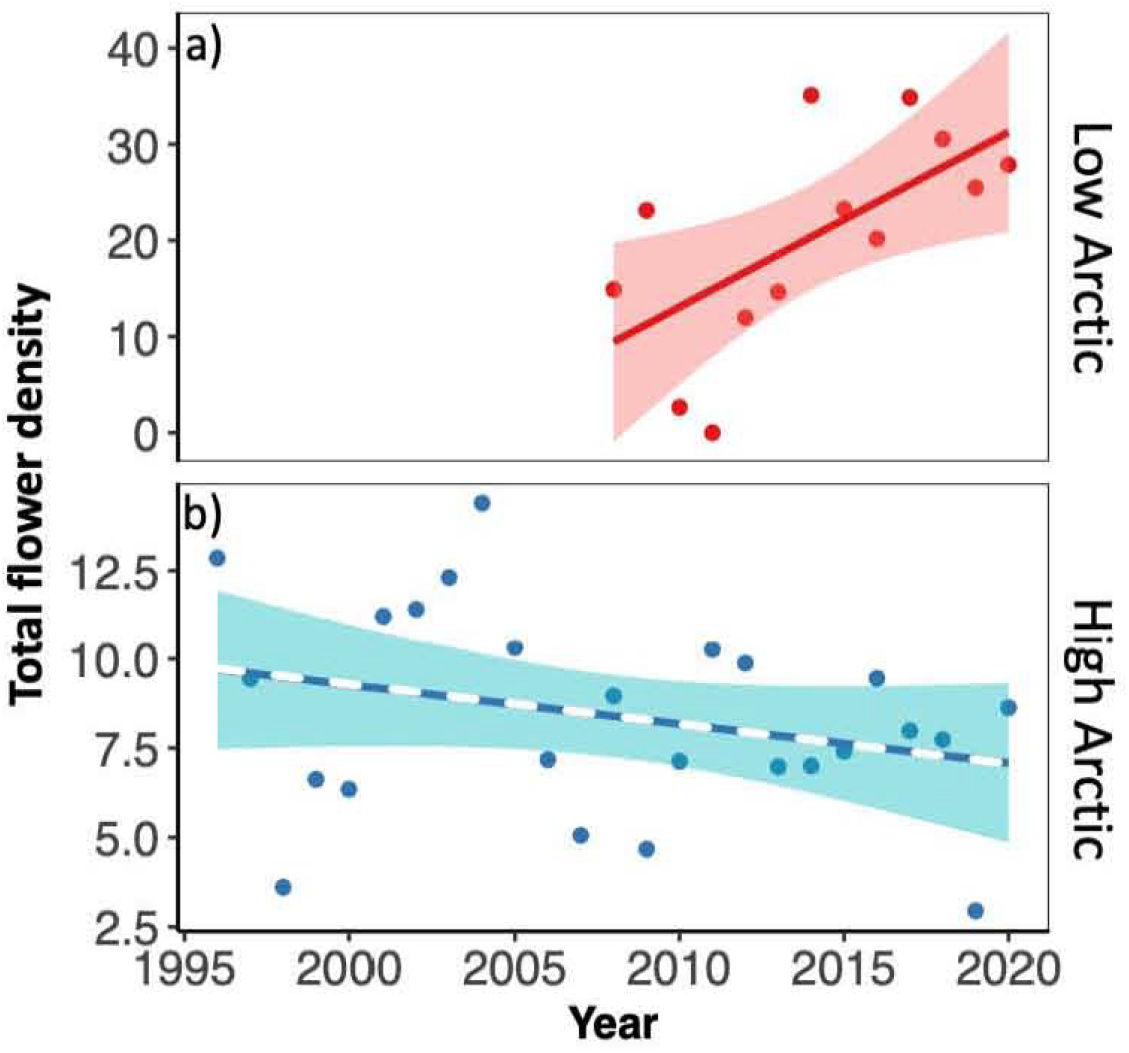
Temporal trends of total flower densities in (a) the low and (b) the high Arctic. Lines show fitted regressions, with the solid line indicating a significant trend, while the dashed line indicates a non-significant effect; coloured bands show the estimated confidence interval of each model. Note the different scales of the y-axis in the two panels, with average flower density being much lower at the high-arctic compared to the low-arctic site.

### Species-level flower production

At both sites, the temporal trends in flower density showed contrasting patterns among species (high Arctic: *Species: Year* F=2.84, p=0.009; low Arctic: *Species: Year* F=8.42, p<0.001). In the low Arctic, we observed a general trend of increasing flower density for *Salix*-female, *Silene* and *Loiseleuria,* compared to *Eriophorum and Salix-male,* which did not show detectable change over time (Fig. 3). In contrast, in the high Arctic, flower density of all species decreased or remained stable over time: *Saxifraga* and *Papaver* showed negative trends, whereas *Cassiope, Dryas, Salix* and *Silene* showed no significant change, with slope estimates being close to zero (Fig. 3).

**Figure 3:**
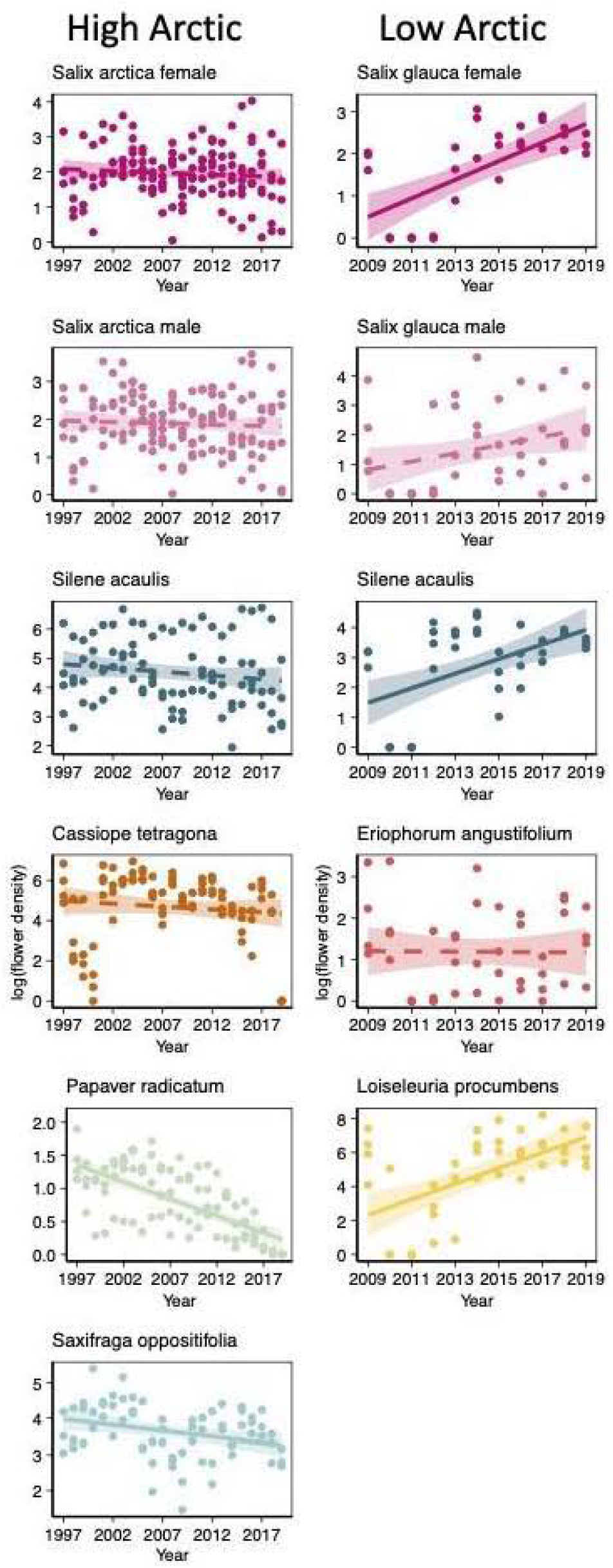
Temporal trends in (log+1) flower density for each species, shown separately for the high- and low-arctic sites. Lines show fitted regressions, with the solid lines indicating significant trends and dashed lines highlighting non-significant models; coloured bands show estimated confidence intervals. Note that the scale of the y-axis varies between panels.

### Drivers of change

Across both sites, the flower density of the previous year (TD) emerged as the most important driver of flower density in the current year. The effect was negative for all nine species, i.e., more flowers in year *t*-1 implied fewer flowers in year *t*. However, this effect differed significantly in magnitude between species (see *Species: TD* in Table 1). The effect of the climatic drivers was highly species-and site-dependent. Only three species experienced a positive effect of summer temperature (ST): *Silene* and *Loiseleuria* in the low Arctic, and *Cassiope* in the high Arctic (see *Species: ST* in Table 1). *Cassiope* in the high Arctic was the only species responding positively to the previous autumn temperature (see *Species: PAT* in Table 1). Finally, we detected a positive effect of snowmelt date for *Loiseleuria* in the low Arctic, and for *Salix-*male and *Cassiope* in the high Arctic. In these species, later snowmelt was reflected in higher flower densities (see *Species: SM* in Table 1).

**Table 1.** Drivers of temporal change in flower density for each species in (a) the high and (b) the low Arctic. Parameter estimates with confidence intervals (95% CI) for the two linear mixed models of the eq. 3. Note: For the effect of previous-year flower density (TD), we show the value of 1-*β_4_*, as reflecting the level of over- or under-compensation. Here, an estimate of 1 implies that flower densities in one year have no effect on flower densities in the next, whereas an estimate below 1 implies that a year of high flower densities is followed by lower densities, and an estimate above ι implies that a year of high flower densities is followed by even higher densities. Marginal R^2^= proportion of variance explained by fixed effects. Conditional R^2^=proportion of variance explained by both fixed and random effects. *CAS=Cassiope tetragona,* SAX=Saxifraga *oppositifolia, SAL=Salix arctica* in the high Arctic, *SAL=Salixglauca* in the low Arctic, *SIL=Silene acaulis, DRY=Dryas integrifolia/octopetala, PAP=Papaver radicatum, LOI=Loiseleuria procumbens.* Green numbers indicate a significant positive while red ones showed a significant negative effect.

|  | a) High Arctic (Zackenberg) |  |  |  | b) Low Arctic (Nuuk) |  |  |  |
| --- | --- | --- | --- | --- | --- | --- | --- | --- |
|  | Estimates | CI | p | df | Estimates | CI | p | df |
| **Species : Summer temperature (ST)** |  |  |  |  |  |  |  |  |
| Species [SAL female] * ST | 0.15 | -0.01 – 0.32 | 0.062 | 427.36 | 0.40 | -0.86 – 1.65 | 0.531 | 52.69 |
| Species [SAL male] * ST | 0.12 | -0.05 – 0.28 | 0.162 | 431.43 | 0.34 | -0.92 – 1.59 | 0.594 | 50.76 |
| Species [SIL] * ST | 0.08 | -0.18 – 0.34 | 0.548 | 427.16 | 1.13 | 0.14 – 2.12 | 0.026 | 49.85 |
| Species [CAS] * ST | 0.47 | 0.27 – 0.68 | <0.001 | 427.16 |  |  |  |  |
| Species [DRY] * ST | 0.12 | -0.05 – 0.28 | 0.180 | 427.16 |  |  |  |  |
| Species [PAP] * ST | 0.04 | -0.17 – 0.24 | 0.731 | 427.16 |  |  |  |  |
| Species [SAX] * ST | -0.03 | -0.43 – 0.38 | 0.900 | 427.16 |  |  |  |  |
| Species [LOI] * ST |  |  |  |  | 1.35 | 0.28 – 2.41 | 0.014 | 49.85 |
| **Species \* Previous autumn temperature (PAT)** |  |  |  |  |  |  |  |  |
| Species [SAL female] * PAT | -0.07 | -0.23 – 0.09 | 0.392 | 427.16 | 1.21 | -0.26 – 2.68 | 0.104 | 52.72 |
| Species [SAL male] * PAT | 0.01 | -0.15 – 0.17 | 0.874 | 431.20 | 1.13 | -0.35 – 2.61 | 0.132 | 52.25 |
| Species [SIL] * PAT | 0.02 | -0.22 – 0.27 | 0.848 | 427.16 | 0.40 | -0.89 – 1.69 | 0.536 | 49.85 |
| Species [CAS] * PAT | 0.19 | 0.00 – 0.38 | 0.050 | 427.16 |  |  |  |  |
| Species [DRY] * PAT | -0.02 | -0.17 – 0.14 | 0.851 | 427.16 |  |  |  |  |
| Species [PAP] * PAT | -0.01 | -0.22 – 0.19 | 0.884 | 427.16 |  |  |  |  |
| Species [SAX] * PAT | 0.01 | -0.36 – 0.38 | 0.948 | 427.16 |  |  |  |  |
| Species [LOI] * PAT |  |  |  |  | 0.62 | -0.51 – 1.75 | 0.274 | 49.85 |
| **Species \* Snowmelt day of the year (SM)** |  |  |  |  |  |  |  |  |
| Species [SAL female] * SM | 0.14 | -0.03 – 0.31 | 0.104 | 427.95 | -0.35 | -1.34 – 0.63 | 0.472 | 51.14 |
| Species [SAL male] * SM | 0.17 | -0.00 – 0.34 | 0.050 | 431.52 | -0.23 | -1.20 – 0.75 | 0.643 | 49.97 |
| Species [SIL] * SM | 0.14 | -0.08 – 0.37 | 0.215 | 427.16 | 0.63 | -0.56 – 1.82 | 0.295 | 49.85 |
| Species [CAS] * SM | 0.35 | 0.10 – 0.60 | 0.006 | 427.16 |  |  |  |  |
| Species [DRY] * SM | 0.05 | -0.12 – 0.22 | 0.549 | 427.16 |  |  |  |  |
| Species [PAP] * SM | -0.01 | -0.24 – 0.23 | 0.990 | 427.16 |  |  |  |  |
| Species [SAX] * SM | 0.12 | -0.35 – 0.58 | 0.622 | 427.16 |  |  |  |  |
| Species [LOI] * SM |  |  |  |  | 1.63 | 0.68 – 2.59 | 0.001 | 49.85 |
| **Species \* Previous year flower density (TD)** |  |  |  |  |  |  |  |  |
| Species [SAL female] * TD | 0.86 | -0.22 – -0.07 | <0.001 | 527.83 | 0.76 | -0.63 – 0.15 | <0.001 | 43.26 |
| Species [SAL male] * TD | 0.86 | -0.22 – -0.07 | <0.001 | 527.69 | 0.61 | -0.79 – 0.01 | <0.001 | 54.51 |
| Species [SIL] * TD | 0.99 | -0.06 – 0.04 | <0.001 | 427.16 | 0.86 | -0.46 – 0.18 | <0.001 | 49.85 |
| Species [CAS] * TD | 0.94 | -0.10 – -0.02 | <0.001 | 427.16 |  |  |  |  |
| Species [DRY] * TD | 0.997 | -0.06 – 0.05 | <0.001 | 427.16 |  |  |  |  |
| Species [PAP] * TD | 0.89 | -0.30 – 0.09 | <0.001 | 427.16 |  |  |  |  |
| Species [SAX] * TD | 0.98 | -0.20 – 0.16 | <0.001 | 427.16 |  |  |  |  |
| Species [LOI] * TD |  |  |  |  | 0.85 | -0.32 – 0.03 | <0.001 | 49.85 |
| **Random Effects** |  |  |  |  |  |  |  |  |
| Residual variance | 0.04 |  |  |  | 0.22 |  |  |  |
| Variance - Plot:Year | 0.72 |  |  |  | 3.09 |  |  |  |
| N - Plot | 490 |  |  |  | 54 |  |  |  |
| N - Year | 28 |  |  |  | 11 |  |  |  |
| Observations | 556 |  |  |  | 76 |  |  |  |
| Marginal R<sup>2</sup> / Conditional R<sup>2</sup> | 0.096 / 0.950 |  |  |  | 0.293 / 0.953 |  |  |  |

At the high-arctic site, the fixed effects of the model including the different drivers (eq. 3) explained ~10% of the variance, while fixed and random effects combined explained ~95% of the variance; the corresponding values for the low-arctic site were ~29% and ~95% (Table 1).

## Discussion

Drawing on unique, long-term data on flower abundance from the Arctic, our analysis revealed pronounced changes in flower density over time. Nonetheless, the more specific temporal trends and the effects of individual climatic drivers proved to differ at both the site-and species levels. Overall, the patterns found were consistent with our *a priori* hypotheses. First, we found substantial differences in recent climatic and flower production trends between sites on opposite sides of Greenland. Second, flower density at the site-level increased over time - but only did so in the low Arctic. Third, we found variation in species trends, with different drivers emerging as important for the flower densities of different species and between sites. Finally, we found a strong trade-off between flower densities in successive years. Below, we will discuss each of these patterns in turn.

### Recent climatic patterns of change differ between a low and a high-arctic site

Current climatic conditions vary substantially between different parts of the Arctic, as do patterns of recent and projected change conditions (Abermann et al. 2017; Kankaanpää et al. 2020; Prevéy et al. 2017; Schmidt et al, 2019b). Consistent with such regional variation, our results show larger climatic changes in northeast Greenland than in southwest Greenland.

While the trends in the annual, summer and autumn temperatures in the low Arctic were not significant, mean annual temperatures changed from negative to positive during the study period (Appendix S4 - Fig. S4a & b). This change in climatic conditions may have substantial direct and indirect impacts, e.g., on plant physiology as well as on water and nutrient availability in the soil. In contrast, in the high Arctic, the mean annual temperatures still remain negative, despite a general trend towards earlier snowmelt dates and higher temperatures (i.e., mean annual, summer and autumn temperature), (Appendix S4 - Fig. S4a & b). The fact that water remains frozen for most of the year maintains strong constraints nutrient and water availability, and on plant metabolism. Such conditions, combined with increased precipitation, have been shown to cause damage to overwintering tissue and flower buds, by exposing plants to detrimental cold winter or spring air temperatures due to a reduced snowpack (Høye et al. 2007a; Inouye 2008; Semenchuk et al. 2013). That such conditions can result in significantly reduced flower density was shown for several species exposed to shallow snowpacks, including *Cassiope tetragona* and with the strongest response observed in *Dryas octopetala* (Høye et al. 2007a, Sememchuk et al, 2020). However, where heavy precipitation in the form of snow prevails, and where wind does not sweep away the snow, a thick layer of snow will provide an effective protective cover at the end of winter (Kelsey et al, 2021, Bokhorst et al, 2011).

### Contrasting temporal trends in flower density between the low- and high-Arctic

With a stronger increase in temperature, we expected stronger trends in flower densities in the high Arctic - assuming that flowering is driven by the same environmental factors as phenology (Krab et al. 2018; Kelsey et al. 2021). Nonetheless, the total density of flowers in the low Arctic nearly doubled over the last 13 years, contrasting with a non-significant (or slowly decreasing) trend in the high Arctic (Fig. 2). Unpacking trends in total flower densities revealed high variability in the direction and magnitude between plant species and sites. For most of the low arctic species, flowering density increased over time, while in the high Arctic, some species (such as *Papaver* and *Cassiope*) showed strong decreasing trends, while others exhibited no significant trend over time (Fig. 3).

### Spatiotemporal variation in the climatic drivers of flower production

In the high Arctic, we found no clear climatic determinant of flower densities. *Cassiope,* one of the later flowering species, proved the species most responsive to the climatic variables: in this species, flower densities increased with summer temperature, the temperature of the previous autumn, and snowmelt date. This finding aligns with our a priori expectations and highlights the need for late-flowering species to accumulate resources during the growing season. At the low-arctic site, both *Silene* and *Loiseleuria* were positively affected by summer temperature, although the former is a late-flowering and the latter is an early-flowering species. Interestingly, *Cassiope* and *Salix* males were the only two species affected by snowmelt timing in the high Arctic. However, in the low Arctic, the early-flowering *Loiseleuria* appeared – as expected – to benefit from late snowmelt. The long-lying snowpack might protect pre-formed buds against spring frosts and provide moisture (Inouye 2008; Niittynen and Luoto 2018; Stewart et al. 2018).

Over time, we detected no significant trend in the environmental driver itself, i.e., the day of snowmelt in the plots where *Loiseleuria* was monitored.

### Trade-off and variation in reproduction investment between years

We found strong support for a trade-off between flowering densities in different years. Specifically, we found a consistent negative effect of the previous year’s flower abundance on the current year’s density, with a stronger effect in the low compared to the high Arctic. This suggests a delayed cost of reproduction, as evidenced by the log-log regression coefficient of flowers in years 1 on densities in year t-1, which were smaller than 1.

Since arctic flowers are typically pre-formed in the previous year (Körner 2003), we might expect a warm autumn to increase the subsequent flower density of early-flowering species, as a longer growth period should allow them to acquire additional resources for bud formation in the autumn. Nonetheless, except for *Cassiope* in the high-arctic site, all species appeared insensitive to such effects. The lack of response to the previous autumn’s temperature seems surprising and might be attributed to a stronger negative effect of the previous year’s flower production and to a lesser extent to current summer conditions.

Overall, the arctic environment is characterised by strong spatio-temporal variation in environmental conditions. In the current data, this was revealed by the high variance associated with the random effects of year and plots (Variance_Year_ & Variance_Plot:Year_, Table 1). This variation can be related to plot variation in micro-topography, affecting moisture (Bannister et al. 2005; Høye et al. 2007a). It suggests that species living in this ecosystem are exposed to highly dynamic and heterogeneous environments. As a result, they need to be flexible and efficient in their resource use, as indicated by increasing species-level trait variation toward colder temperatures (Siefert et al. 2015).

### Implications of changes in flower production

Our analyses reveal substantial changes in both total and species-specific flower densities over time. At the low-arctic site, we observed an increase in overall flowering densities, in contrast to a downward trend in flowering density at the low-arctic site driven by Saxifraga and *Papaver.* For pollinators and other plant-associated taxonomic groups, a decrease in the number of flowers may increase competition for access to pollen, nectar or seeds. In a resource-limited ecosystem such as the Arctic, this resource availability change can directly affect diversity and community structure. For plants, an increase in flower production could, for example, directly affect competition for pollinators (Hocking 1968; Tiusanen et al. 2020). Adding to this are changes in the relative phenology of plants, which may either increase or decrease competition, depending on how changes affect the temporal overlap between more and less-attractive plant species (Tiusanen et al. 2020). Overall, a potential future decrease in flower production, as observed for *Saxifraga* and *Papaver* in the high Arctic, has the potential to affect the long-term recruitment of new individuals (Inouye 2008) and long-distance dispersal. However, since many tundra plant species may combine sexual reproduction with a clonal and vegetative strategy, it is difficult to directly link our results to the population dynamics of all nine species (Wipf 2010). Further studies on mortality, colonization and vegetative growth rates are needed to connect the current patterns to plant population dynamics at the ecosystem scale. Our current findings suffice to suggest major changes in the flowering of arctic plants, with large variation between species and sites.

### Conclusions

In this study, we used long-term time series of flower abundance of nine species of arctic tundra to disentangle the effects of climatic change and reproductive cost on flower density over space and time. We found that the temporal trends and effects of past and current conditions on flower density differed at both the site and species levels. This heterogeneity of site- and species-level responses highlights the complex nature of vegetation–climate interactions in arctic tundra plant communities. It suggests that local communities, as well as co-occurring species, may exhibit contrasting responses to climate change, as they respond to different specific climatic drivers. These asymmetric patterns may affect vegetation dynamics, as well as lead to cascading effects to other trophic levels. Per extension, they may reverberate across the tundra ecosystem.

## Acknowledgements

All data for this study were obtained from the Greenland Ecosystem Monitoring database (https://data.g-e-m.dk). We are indebted to all collaborators and field assistants at Zackenberg and Nuuk over the years. We thank Bastien Parisy, Helena Wirta, Sonia Saine and Michael Belluau for stimulating discussions.

## Ethics approval

NA

## Consent to participate

NA

## Consent for publication

NA

## Availability of data and material

All data are freely available from the Greenland Ecosystem Monitoring Programme, GEM, https://data.g-e-m.dk.

## Code availability

https://github.com/AntoineBeckerScarpitta/Zackenberg_Flowering

## Authors’ contributions

TR, NMS, ABS, and LHA conceived the project. ABS and LHA prepared and analysed the data in close collaboration with GB, TR & NMS. NMS and KR coordinated the data collection (at Zackenberg and Nuuk research stations respectively). ABS, LHA wrote the first draft of the manuscript, with substantial input from TR and NMS. All authors then contributed to later versions of the manuscript.

## List of appendices

Appendix S1 - Map of the study site

Appendix S2 - Flower abundance time series sampling design

Appendix S3 - Study species

Appendix S4: Climatic trends of Precipitation (mm), Temperature (°C) and Snowmelt day:

- Figure S4a: Mean annual trends for Precipitation and Temperature.
- Figure S4b: Mean seasonal trends for Precipitation and Temperature
- Table S4b: Temporal trends for the mean seasonal Temperature and Precipitation in the low-and high-arctic sites
- Figure S4c: Temporal trends for the Snowmelt day of the year
- Table S4c: Temporal trends for the Snowmelt day of the year in the low-and high-arctic sites.

