## Supplements for "Diverging trends and drivers of Arctic flower production over space and time"

**Appendix S1** - **Map of the study sites:** (a) the low-arctic site (Kobbefjord research station, Raundrup, Olsen, et al. 2020) and (b) the high-arctic site (Zackenberg research station, Schmidt, Hansen, et al. 2019a). Dots within landscapes indicate the permanent monitoring plots where flower densities were recorded, with abbreviations identifying the focal plant species: CAS=*Cassiope tetragona,* DRY=*Dryas integrifolia/octopetala*, ERI=*Eriophorum angustifolium*, LOI=*Loiseleuria procumbens*, PAP=*Papaver radicatum,* SAX=*Saxifraga oppositifolia*, SAL=*Salix sp.* (with the species monitored being *Salix glauca* at Nuuk and *Salix arctica* at Zackenberg)*,* SIL=*Silene acaulis*.

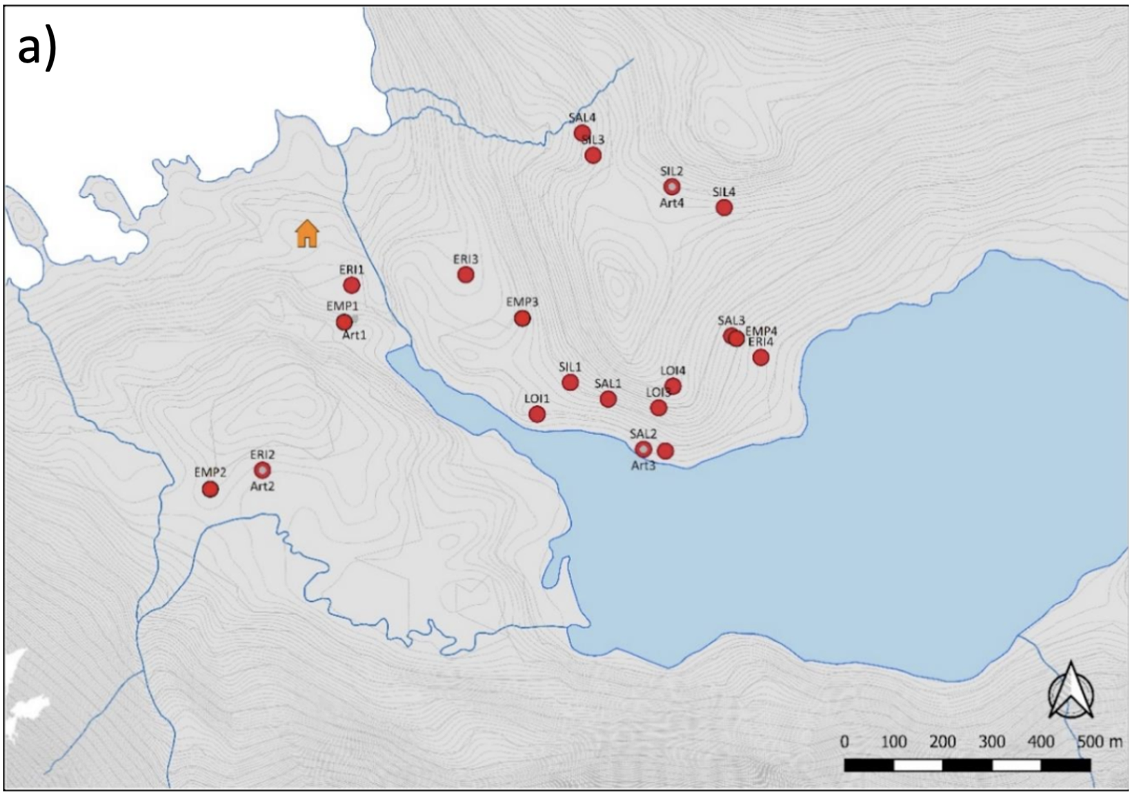

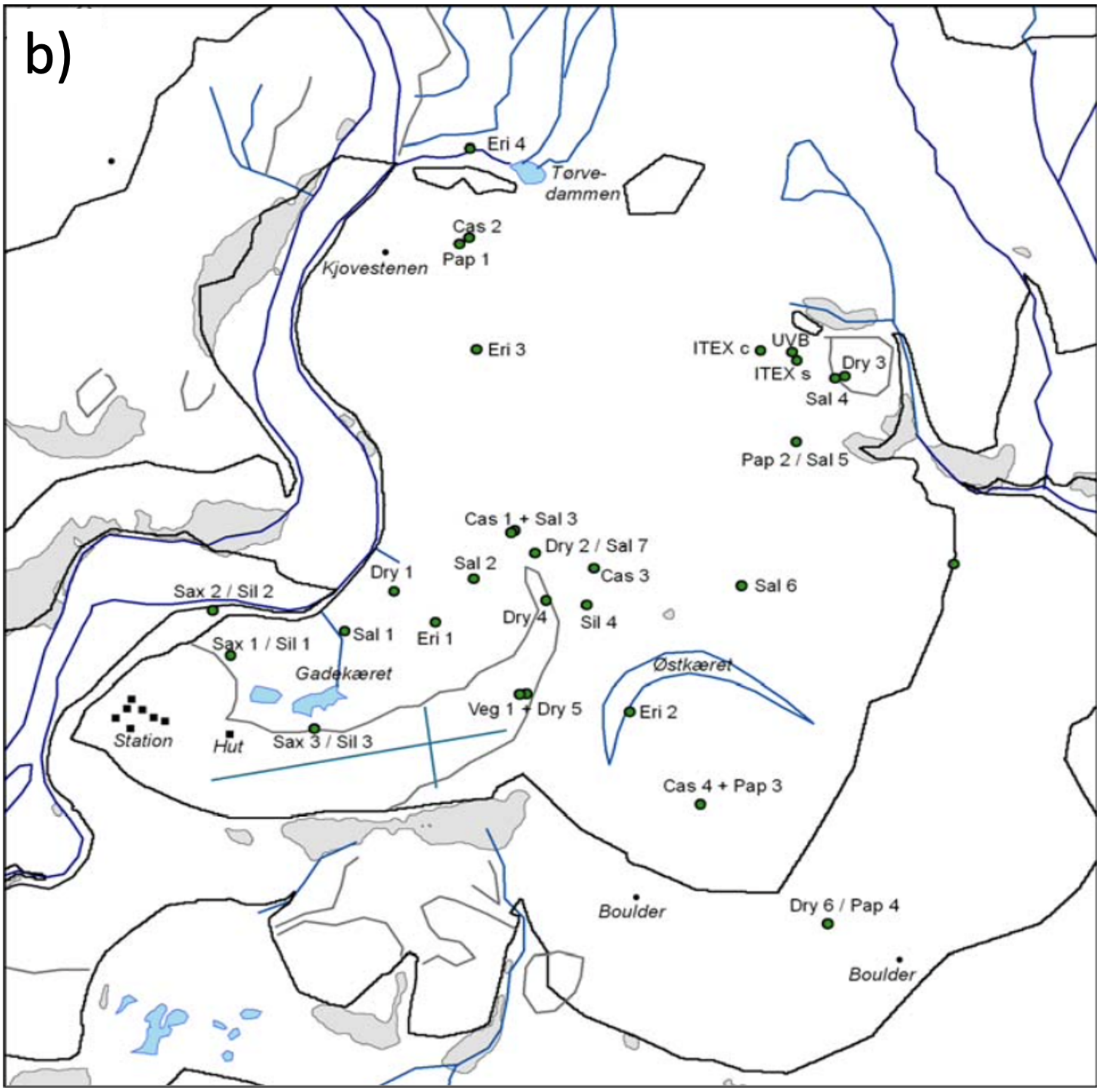
b)
Eri 4
Tørve-
dammen
Cas 2
Pap 1
Kjovestenen
Eri 3
ITEX c
UVB
ITEX s
Dry 3
Sal 4
Pap 2 / Sal 5
Cas 1 + Sal 3
Dry 2 / Sal 7
Cas 3
Sal 6
Sal 2
Dry 1
Sax 2 / Sil 2
Sal 1
Eri 1
Dry 4
Sil 4
Sax 1 / Sil 1
Gadekæret
Østkæret
Veg 1 + Dry 5
Eri 2
Station
Hut
Sax 3 / Sil 3
Cas 4 + Pap 3
Boulder
Dry 6 / Pap 4
Boulder

**Appendix S2 – Flower abundance time series sampling design**, showing the number of plots in the two study sites for each species and each year. Species names as in Appendix 1; the high Arctic - Zackenberg with a duration of 25 years, and the low Arctic - Nuuk of 13 years.

| | Zackenberg | | | | | | | Nuuk | | | | |
|---|---|---|---|---|---|---|---|---|---|---|---|---|
| | CAS | DRY | PAP | SAL_female | SAL_male | SAX | SIL | ERI | LOI | SAL_female | SAL_male | SIL |
| 1996 | 4 | 6 | 4 | 4 | 4 | 3 | 4 | - | - | - | - | - |
| 1997 | 4 | 6 | 4 | 4 | 4 | 3 | 4 | - | - | - | - | - |
| 1998 | 4 | 6 | 4 | 4 | 4 | 3 | 4 | - | - | - | - | - |
| 1999 | 4 | 6 | 4 | 4 | 4 | 3 | 4 | - | - | - | - | - |
| 2000 | 4 | 6 | 4 | 4 | 4 | 3 | 4 | - | - | - | - | - |
| 2001 | 4 | 6 | 4 | 4 | 4 | 3 | 4 | - | - | - | - | - |
| 2002 | 4 | 6 | 4 | 4 | 4 | 3 | 4 | - | - | - | - | - |
| 2003 | 4 | 6 | 4 | 7 | 6 | 3 | 4 | - | - | - | - | - |
| 2004 | 4 | 6 | 4 | 7 | 7 | 3 | 4 | - | - | - | - | - |
| 2005 | 4 | 6 | 4 | 7 | 7 | 3 | 4 | - | - | - | - | - |
| 2006 | 4 | 6 | 4 | 7 | 7 | 3 | 4 | - | - | - | - | - |
| 2007 | 4 | 6 | 4 | 7 | 7 | 3 | 4 | - | - | - | - | - |
| 2008 | 4 | 6 | 4 | 7 | 7 | 3 | 4 | 4 | 2 | 3 | 4 | 2 |
| 2009 | 4 | 6 | 4 | 7 | 7 | 3 | 4 | 4 | 4 | 3 | 4 | 3 |
| 2010 | 4 | 6 | 4 | 7 | 7 | 3 | 4 | 4 | 4 | 3 | 4 | 4 |
| 2011 | 4 | 6 | 4 | 7 | 7 | 3 | 4 | 4 | 4 | 3 | 4 | 4 |
| 2012 | 4 | 6 | 4 | 7 | 7 | 3 | 4 | 4 | 4 | 3 | 4 | 4 |
| 2013 | 4 | 6 | 4 | 6 | 6 | 3 | 4 | 4 | 3 | 3 | 4 | 4 |
| 2014 | 4 | 6 | 4 | 7 | 7 | 3 | 4 | 4 | 4 | 3 | 4 | 4 |
| 2015 | 4 | 6 | 4 | 7 | 7 | 3 | 4 | 4 | 4 | 3 | 4 | 4 |
| 2016 | 4 | 6 | 4 | 7 | 7 | 3 | 4 | 4 | 4 | 3 | 4 | 4 |
| 2017 | 4 | 6 | 4 | 6 | 6 | 3 | 4 | 4 | 4 | 3 | 4 | 4 |
| 2018 | 4 | 6 | 4 | 6 | 5 | 3 | 4 | 4 | 4 | 3 | 4 | 4 |
| 2019 | 4 | 6 | 4 | 4 | 5 | 3 | 4 | 4 | 4 | 3 | 4 | 4 |
| 2020 | 4 | 6 | 4 | 6 | 6 | 3 | 4 | 4 | 4 | 3 | 4 | 4 |

### Appendix S3 – Ecological details on the species studied.

Due to the advanced development of buds at the end of the previous growing season, *Saxifraga oppositifolia* L. is the earliest species to flower during spring in May-June (Stenström, et al 1997). It is a densely tufted succulent forb which is common on varying types of soil.

Species flowering in June-July are: *Loiseleuria procumbens* (L.) Desvaux, *Salix arctica* L., *Salix glauca* L., *Eriophorum angustifolium* L., and *Papaver radicatum* Rottb. Of these, *Loiseleuria procumbens* is an evergreen prostrate dwarf-shrub, growing on acid and dry heaths. *Salix arctica* is a short-statured shrub with a large ecological niche widely distributed across the Arctic heath and fell fields. *Salix glauca* is a deciduous shrub that typically grows on sand and cobbles among granitic boulders, sandy alluvium or scree slopes. *Eriophorum angustifolium* is a perennial graminoid, commonly found on peaty wet soil; and *Papaver radicatum* is a hardy forb forming tufts. It is a generalist species widely distributed across the Arctic fell fields.

*Cassiope tetragona* (L.) D. Don, *Silene acaulis* (L.) Jacq are the latest species to flower in July-August. Of these, *Cassiope tetragona* is an evergreen dwarf shrub with a large ecological niche, *Silene acaulis* growing in compact cushions is a forb occurring on sandy open fell fields and heath.

Finally, the hybrid *Dryas integrifolia/octopetala* is a cushion-forming evergreen shrub, very common in non-acidic arctic tundra heath and has a very broad flowering niche from June to August.

**Appendix S4 - Climatic trends of Precipitation (mm), Temperature (°C) and Snowmelt day** at the low-arctic site for the period 2007-2020 and the high-arctic sites for the period 1995-2020.

**Figure S4a: Mean annual trends for Precipitation and Temperature.** Mean annual precipitation increased in the high Arctic ($\beta$=0.001±0.0003, t=3.42, p=0.002), while it decreased in the low Arctic ($\beta$= -0.003±0.002, t=-1.93, p=0.04). We detected significant warming in the high Arctic ($\beta$=0.067±0.02, t=3.52, p=0.001), while no change in annual temperature was detected in the low Arctic ($\beta$=0.16±0.14, t=1.12, p=0.3).

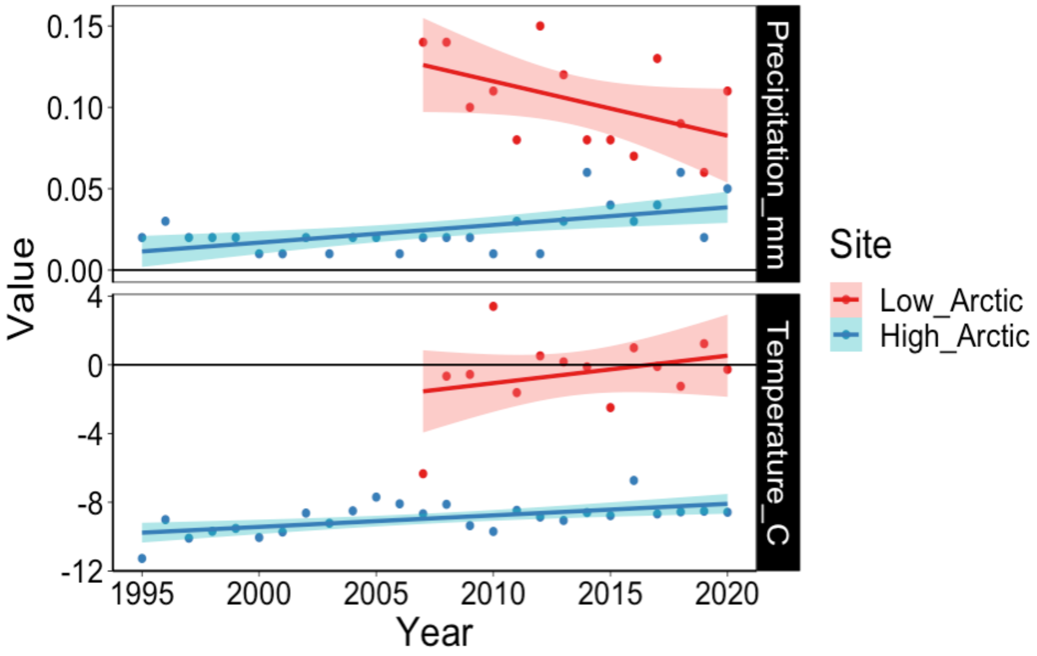

**<u>Figure S4b:</u> Mean seasonal trends for Precipitation and Temperature,** with *Summer*=June to August, and *Autumn*=September to November; trend estimates are given in Table S4b.

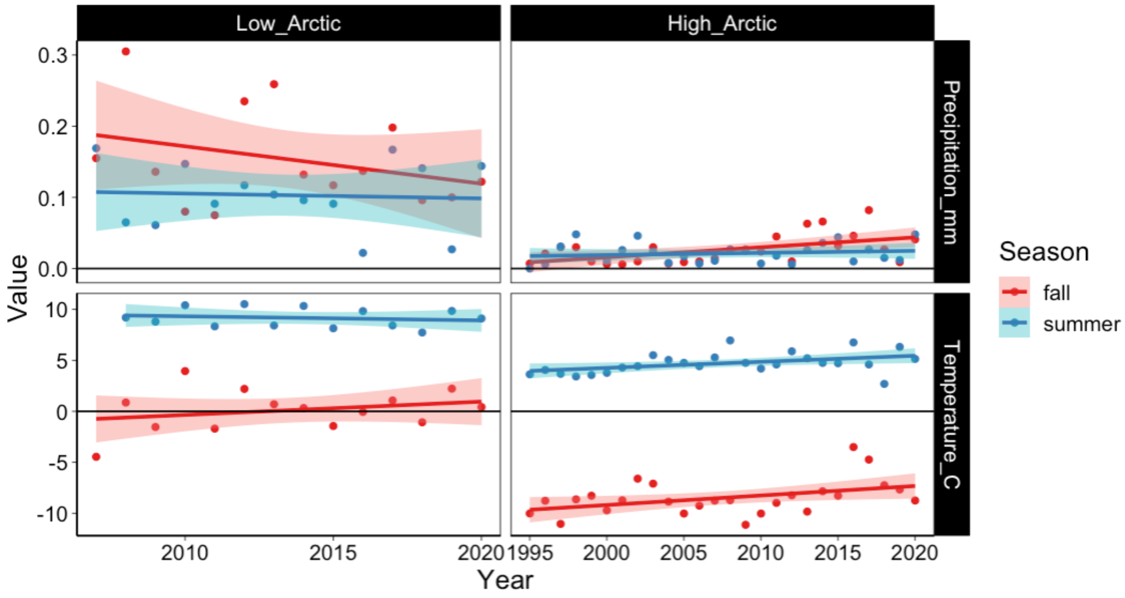


<u>Table S4b</u>: **Temporal trends in mean seasonal temperature and precipitation** in the high- (a) and low- (b) arctic sites. Bold values are significant.

| | Estimate | std.error | t | p |
|---|---|---|---|---|
| **a) High Arctic** | | | | |
| **Precipitation** | | | | |
| (Intercept) | -1.67 | 0.615 | -2.72 | **0.009** |
| Year:Season_fall | 0.001 | 0.0003 | 2.76 | **0.008** |
| Year:Season_summer | 0.001 | 0.0003 | 2.75 | **0.008** |
| **Temperature** | | | | |
| (Intercept) | -155.5 | 47.9 | -3.25 | **0.002** |
| Year:Season_fall | 0.073 | 0.024 | 3.07 | **0.003** |
| Year:Season_summer | 0.08 | 0.024 | 3.35 | **0.001** |
| **b) Low Arctic** | | | | |
| **Precipitation** | | | | |
| (Intercept) | 6.11 | 5.65 | 1.08 | 0.3 |
| Year:Season_fall | -0.003 | 0.003 | -1.06 | 0.3 |
| Year:Season_summer | -0.003 | 0.003 | -1.06 | 0.3 |
| **Temperature** | | | | |
| (Intercept) | -105.43 | 193 | -0.635 | 0.5 |
| Year:Season_fall | 0.052 | 0.082 | 0.640 | 0.5 |
| Year:Season_summer | 0.057 | 0.082 | 0.695 | 0.5 |

**<u>Figure S4c</u>: Temporal trends of Snowmelt day of the year** in the low- (left) and high-arctic (right) sites. The snowmelt day of the year was calculated as the date when 50% of the plot reached 50% of snow cover. No significant trends were found in the low Arctic, while all plots (each species was inventoried in an independent set of plots, Appendix S1) showed a significant decrease in the snowmelt day in the high Arctic, meaning earlier melting of snow; trend estimates are given in Table S4c.

DOY-120=30 April; DOY-140=20 May; DOY-160=9 June; DOY-180=29 June.

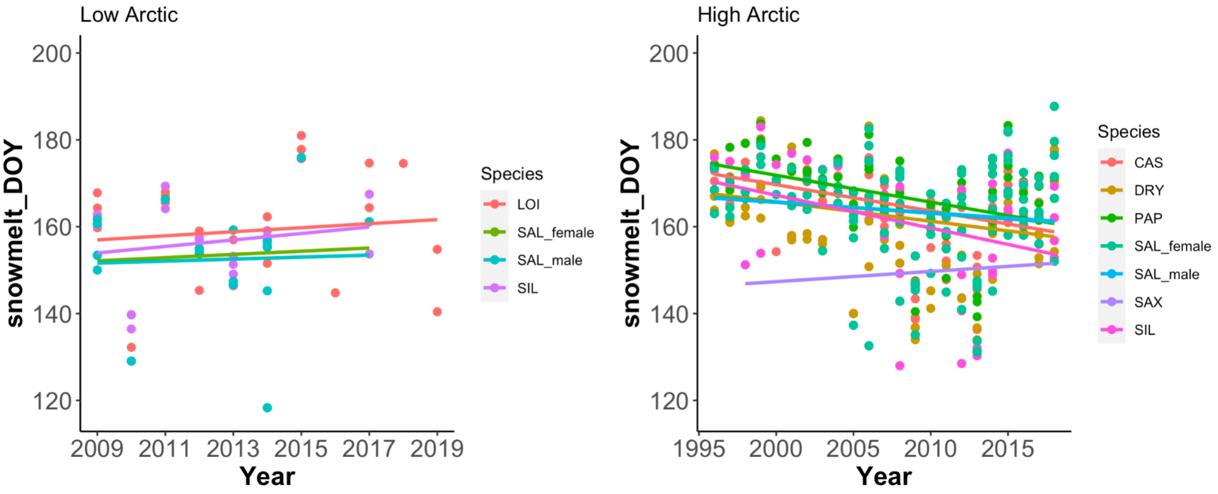


Table S4c: Temporal trends of Snowmelt day of the year for each species plots in the high- (a) and low- (b) arctic sites. Bold values are significant.

| | Estimate | std.error | t | p |
|---|---|---|---|---|
| **a) High Arctic** | | | | |
| Year:Species_CAS | -0.603 | 0.141 | -4.28 | **<0.001** |
| Year:Species_DRY | -0.447 | 0.182 | -2.45 | **0.01** |
| Year:Species_PAP | -0.619 | 0.147 | -4.21 | **<0.001** |
| Year:Species_SAL_female | -0.260 | 0.177 | -1.47 | 0.1 |
| Year:Species_SAL_male | -0.240 | 0.179 | -1.34 | 0.1 |
| Year:Species_SAX | 0.235 | 0.476 | 0.493 | 0.6 |
| Year:Species_SIL | -0.755 | 0.309 | -2.44 | **0.01** |
| **b) Low Arctic** | | | | |
| Year:Species_LOI | 0.466 | 0.875 | 0.544 | 0.592 |
| Year:Species_SAL_female | 0.369 | 1.65 | 0.224 | 0.826 |
| Year:Species_SAL_male | 0.230 | 1.39 | 0.165 | 0.871 |
| Year:Species_SIL | 0.747 | 0.859 | 0.869 | 0.395 |
